# Enhanced intracranial aneurysm development in a rat model of polycystic kidney disease

**DOI:** 10.1101/2024.02.28.582650

**Authors:** Anne F. Cayron, Sandrine Morel, Maral Azam, Julien Haemmerli, Tomohiro Aoki, Philippe Bijlenga, Eric Allémann, Brenda R. Kwak

## Abstract

**Background:** Polycystic kidney disease (PKD) patients have a high intracranial aneurysms (IAs) incidence and risk of rupture. The mechanisms that make PKD patients more vulnerable to IA disease are still not completely understood. The PCK rat is a well-known PKD model and has been extensively used to study cyst development and kidney damage. Here, we used this rat model to study IA induction and vulnerability.

**Methods:** IAs were induced in wild-type (WT) and PCK rats and their incidence was followed. Variation in the anatomy of the circle of Willis was studied in PCK rats and PKD patients. The expression of tight junction proteins was examined by immunohistochemistry in rats and in human ruptured and unruptured IAs from patients enrolled in the @neurIST study.

**Results:** An increased frequency of fatal aortic dissection was unexpectedly observed in PCK rats after IA induction, which was due to modifications in the elastic architecture of the aorta in combination with the induced hypertension. Interestingly, IAs developed faster in PCK rats compared to WT rats. Variations in the anatomy of the circle of Willis were identified in PCK rats and PKD patients, a risk factor that may (in part) explain the higher IA incidence found in these groups. At 2-weeks after induction, the endothelium of IAs from PCK rats showed a decrease in the tight junction proteins zonula occludens-1 and claudin-5. This decrease in tight junction proteins was also observed in the endothelium of human ruptured IAs compared to unruptured IAs, making it a potential marker of IA wall vulnerability.

**Conclusions:** Our study showed that PCK rats are more sensitive to IA induction. Variations in the anatomy of the circle of Willis and impaired regulation of tight junction proteins might put PCK rats and PKD patients more at risk of developing vulnerable IAs.

## INTRODUCTION

Intracranial aneurysms (IAs) are abnormal enlargements of the lumen of arteries in the circle of Willis (CoW), which affects 2 to 5% of the population^1,2^. IA rupture leading to subarachnoid hemorrhage (SAH) is fatal in 25 to 50% of the cases^3^. Unruptured IAs can be preventively treated, but these interventions are rather perilous with ∼4,2% risk of mortality and ∼3,6% risk of morbidity^4^. Thus, it is essential to identify unstable IAs prone to rupture accurately.

IAs typically develop at bifurcations within the CoW, an arterial network responsible for supplying blood to the brain^5^. The CoW is composed of the anterior cerebral arteries (ACA), the anterior communicating artery (ACOM), the internal carotid arteries (ICA), the middle cerebral arteries (MCA), the posterior communicating arteries (PCOM), the posterior cerebral arteries (PCA) and the basilar artery (BA) and is divided in the anterior and posterior circulations fed by the ICAs and BA, respectively^6^. Many morphological variations in the CoW have been found in the general population^7^. Those variations modify the hemodynamics and the wall shear stress applied to endothelial cells (ECs)^8^. Wall shear stress is known to have a critical role in the initiation of IAs^5^, and CoW variations have been correlated with this process^9,10^.

Polycystic kidney disease (PKD) is a genetic disease characterized by an excessive development of renal cysts, which deteriorates kidney function and may lead to organ failure^11^. Interestingly, PKD has been associated with a high IA incidence and risk of rupture^11,12^. PKD patients carry a mutation in the PKD1, PKD2 or PKHD1 genes affecting the presence or the function of primary cilia^11^. In ECs lining the lumen of the arterial wall, primary cilia act as mechano-sensors of blood flow^5^. We have recently shown *in vitro* that primary cilia dampen the endothelial response to an aneurysmal pattern of disturbed flow^13^. The tight junction (TJ) protein Zonula Occludens-1 (ZO-1) was identified as a central regulator of primary cilia-dependent junction integrity^13^. As TJs control endothelial permeability, dysfunctional endothelial TJs have been associated with multiple vascular diseases^14^. Claudin-5 (CLDN5) is a TJ protein highly expressed in intracranial endothelium and plays a key role in maintaining the blood brain barrier^15,16^. We hypothesize that a decrease of TJ proteins might explain why PKD patients develop IAs more frequently and these IAs are more at risk to rupture.

In this study, PCK rats were used as model of PKD and compared to wild-type (WT) rats. The PCK rat, which carries non-functional primary cilia^17^, was first described in the early 21^st^ century^18,19^ and has been extensively used to study cyst development and hepatorenal damage^17^. However, PCK rats were never used to study IA development. We found that IAs developed faster in PCK rats compared to WT rats. Moreover, variations in the anatomy of the CoW were identified in PCK rats and PKD patients, a risk factor that may (in part) explain the high IA incidence found in these groups. At 2-weeks after induction, the endothelium of IAs from PCK rats showed a lower content in TJ proteins ZO-1 and CLDN5, compared to WT rats. This reduced TJ protein expression was also observed in the endothelium of human ruptured IAs compared to unruptured IAs of patients enrolled in the @neurIST study.

## METHODS

### Data Availability

Data supporting the findings of this study are available from the corresponding author upon reasonable request. All research materials listed in the Methods section are included in the Major Resource Tables available in Supplemental Material.

### Rat IA model

The experiments were approved by the Swiss federal and cantonal veterinary authorities (license GE21519A) and performed according to the Guide for the Care and Use of Laboratory Animals and the Swiss national animal protection laws. The animal experiments complied with the National Institute of Health’s Guide for the Care and Use of Laboratory Animals and the Animal Research Reporting *In Vivo* Experiments (ARRIVE) guidelines.

Five to 9 weeks old male WT and PCK (PCK/CrljCrl-Pkhd1^pck^/CRL) CD Sprague-Dawley rats were obtained from Charles River Laboratories. The animals were randomly selected and placed in cages (2-5 animals per cage). Animals were marked with tail marks and the acclimatization phase lasted at least 7 days. Rats were held in an enriched environment with *ad libitum* access to water and food. After acclimatization, rats were accommodated to the restrainer chamber to measure systolic blood pressure for 3 consecutive days. The protocol to induce IAs was applied to 30 WT and 33 PCK rats. Eight WT and 12 PCK controls rats did not undergo any surgery, were fed with a normal diet and were killed in parallel with rats exposed to the IA induction protocol. All animals were monitored and daily scored for general well-being during the first 3 days post-surgery and then twice a week until the end of the experiment.

Following the well-known protocol of Aoki *et al.*^20,21^, IAs were induced at the right olfactory artery (OA) and ACA bifurcation. Briefly, under general anesthesia with inhalation of isofluorane (4% for induction, 2% for maintenance), hemodynamic stress was increased on the right half of the CoW by ligation of the left common carotid artery (LCCA). Hypertension was induced by ligation of the left renal artery in combination with a high salt diet (8%). The arterial wall was weakened by addition of 0.12% β-aminopropionitrile (BAPN) to the diet. To reduce pain, two subcutaneous injections of buprenorphine (0.05 mg/kg in 100 μL) were given 30 min before surgery and 4-6 hours later. Buprenorphine was also added to the drinking water (0.05 mg/kg) for 2 days post-surgery. Systolic blood pressure was regularly measured by the tail-cuff method. Animals were killed at 1- to 8-weeks post-surgery. Any animal that died during the experiment was meticulously examined to determine the cause of death.

### Clinical and radiological data

Patients have been recruited at the Geneva University Hospitals. The inclusion criteria were as follows: 1) IA identified on the basis of angiographic appearance (3D-Digital Subtraction Angiography, 3D-Magnetic Resonance Angiogram, or 3D-Computed Tomography Angiogram) as well as availability of surgical documentation; 2) age older than 18 years; 3) patient provided informed consent. The exclusion criteria were as follows: 1) lack of angiographically proven IA on 3D-Digital Subtraction Angiography, 3D-Magnetic Resonance Angiogram or 3D-Computed Tomography Angiogram; 2) failure to contribute to clinical data; and 3) refusal to provide informed consent. This study is in accordance with the Helsinki Declaration of the World Medical Association and was approved by the Geneva State Ethics Commission for Research as part of the @neurIST study (PB 2022-00426 previously PB_2018-0073 and 07-056). All patients approved for the use of their data and biological samples in the field of cerebrovascular research.

For the analysis of CoW organization, magnetic resonance images (MRIs) of 16 PKD patients were retrieved from the @neurIST cohort. MRIs of 16 matched non-PKD patients were also selected from the @neurIST cohort such that there were no differences between the two groups in terms of sex, positive family history for IAs and IA multiplicity. For the purpose of the analysis, a time of flight (TOF) sequence was used. CoW were classified by 3 blinded observers using the TOF images according to the classification presented in **Figure S1** which was adapted from Kızılgöz *et al*^6^.

### Human IA samples

For histological studies, 38 unruptured and 19 ruptured human IA domes localized at the MCA in non-PKD patients were retrieved from the Swiss AneuX biobank^22^. Human saccular IAs samples were obtained and processed as previously described^22^.

### Rat samples

Rats were euthanized under anesthesia with an intraperitoneal injection of ketarom (ketasol 100 mg/kg and xylazine 10 mg/kg) and transcardially perfused with 4% paraformaldehyde. The CoW, aortic arch and right kidney were collected from all rats, fixed overnight in 4% paraformaldehyde and transferred to a 30% sucrose-phosphate-buffered saline solution until complete dehydration. Prior to sample embedding, the complete CoW, right OA-ACA bifurcation and aortic arches of all rats were imaged using a Stemi 508 stereo microscope (Carl Zeiss). Right kidneys were weighed. Right OA-ACA bifurcation, aortic arch and right kidney of all animals were embedded in OCT compound, snap frozen and stored at -80°C.

Images of the CoW of all WT (n=38) and PCK (n=44) rats were analyzed using the ImageView software (ImageView v4.11.18709.20210403) to study their anatomical organization. The CoW of one PCK rat that died prematurely was too damaged and was not studied. Aortic arches of animals that died prematurely were studied to find potential anomalies that could have caused the death of the animal.

### Histology, Immunohistochemistry and immunofluorescence

Human IA samples were sectioned at 5 μm and conserved at 4°C. Immunoreactivity was retrieved by pressure cooker treatment (3 min) in citrate buffer. ECs were immunolabeled for CD31 (#DLN-33600, Dianova, 1/25), ZO-1 (#61-7300, Invitrogen, 1/100) and CLDN5 (#PAB27037, Abnova, 1/1000). Labelings were visualized using secondary antibody coupled to streptavidin-biotin peroxidase complex and 3,3’-diamino-benzidine chromophore using the EnVision Flex system (#K8023, DAKO) or the VECTASTAIN Kit (#PK-7800, Vector Laboratory) combined with ImmPACT DAB Substrate Kit (#SK-4105, Vector Laboratory). Hemalun (#1.09249, Merck) was used as counterstaining. Samples were mounted with Aquatex (#108562, Sigma Aldrich). Stainings were performed on consecutive sections.

The right OA-ACA bifurcations, aortic arches and right kidneys of rats were sectioned at 6 μm and conserved at -20°C. Kidney sections were stained with hematoxylin/eosin. Aortic arch sections were stained for Martius Scarlet blue to visualize fibrin and Victoria blue to study the elastic architecture. Right OA-ACA bifurcation sections were fixed in 4% paraformaldehyde for 15 min, permeabilized in ice-cold 100% methanol for 3 min and blocked with 2% bovine serum albumin (#A1391, AppliChem) for 15 min. ECs were immunolabeled for CD31 (#DLN-33600, Dianova, 1/100), ZO-1 (#61-7300, Invitrogen, 1/100) and CLDN5 (#PAB27037, Abnova, 1/200). A FITC-conjugated goat anti-rabbit antibody (#711-095-152, Jackson Lab, 1/100) was used for signal detection. Nuclei and internal elastic lamina were counterstained with 4’,6-diamidino-2-phenylindol (DAPI) and 0.003% Evans Blue, respectively. Samples were mounted with Vectashield antifade mounting medium (#H-1000-10, Vector Laboratories). Stainings were performed on consecutive sections.

### Image analysis

Stainings on human IA sections, rat aortic arches and kidneys were scanned at 10X magnification using the Axio Scan.Z1 automated slide scanner (Carl Zeiss) and images were processed by a blinded observer using the Zen 2 software (Version 3.4.91, Carl Zeiss). Using the ImageJ (FIJI) software (Java 1.8.0, NIH), CD31, ZO-1 and CLDN5 staining on human IA sections was quantified by a blinded observer at 5 different locations along the endothelium using previously established methods^13^. Results were expressed as the mean of the 5 measurements for each sample. Fluorescent stainings on rat right OA-ACA bifurcation sections were imaged using the epifluorescent Zeiss Axiocam Imager Z1 (Carl Zeiss) equipped with an AxioCam 506 mono camera (Carl Zeiss). Images were processed and analyzed using the Zen 2 software (Version 3.4.91, Carl Zeiss) by a blinded observer. Similar to the human sections, CD31, ZO-1 and CLDN5 stainings were quantified by a blinded observer at 5 different locations along the endothelium at the level of the right OA-ACA bifurcation, where the IA developed, and in the ACA. Results were expressed as the mean of the 5 measurements for each sample.

### Statistical analysis

Statistical analysis was performed using GraphPad Prism (Version 10.1.1, GraphPad). The distribution of the data was assessed by the D’Agostino-Pearson Normality test. Normally distributed data are expressed as mean±SD. Data with non-normally distributed data (including n<8) are expressed as median ± interquartile range. The presence of outliers was identified using the ROUT method (Q=1%) and removed from the statistical analysis if present. The difference between the means of 2 independent groups with normally distributed data was analyzed using a one- or two-tailed Student’s *t*-test. One-tailed *t*-test was used when a difference in a specific direction was expected. The difference between the median of 2 independent groups with non-normally distributed data was analyzed using a Mann-Whitney U-test. Difference between the median of multiple groups with non-normally distributed data was analyzed using a Kruskal-Wallis test and Dunn’s multiple comparisons test. Comparison of distributions have been performed using the Fisher’s exact test. P<0.05 indicates statistical significance: *P<0.05, **P <0.01, ***P <0.001 and ****P <0.0001.

## RESULTS

### WT and PCK rats display similar increases in systolic blood pressure

To induce IAs, young (6-10 weeks) WT and PCK rats were exposed to ligation of the LCCA and the left renal artery followed by a high salt diet enriched with BAPN. The bodyweight of WT and PCK rats were similar (∼320 gram) at the time of surgery and although PCK rats seemed to gain less weight than WT rats these differences were not significant during the full experiment (**Figure 1A**). As expected, hematoxylin/eosin staining on the right kidney revealed the presence of cysts in PCK rats (**Figure 1B**) which were absent in WT rats (**Figure 1C**). Likewise, we observed increased weight of the right kidney in control PCK rats (**Figure 1D**) and in PCK rats after surgery **(Figure 1E**) compared to control WT rats and WT rats after surgery. Finally, the systolic blood pressure was carefully monitored before and after surgery. In both WT and PCK rats, the systolic blood pressure raised in 1 week by 30 mmHg reaching ∼160 mmHg after which it remained elevated, thus confirming the induced hypertension in WT and PCK rats in response to renal artery ligation and high salt diet (**Figure 1F**). Despite the polycystic kidneys of PCK rats, no differences in blood pressure were observed between WT and PCK rats.

**Figure 1:**
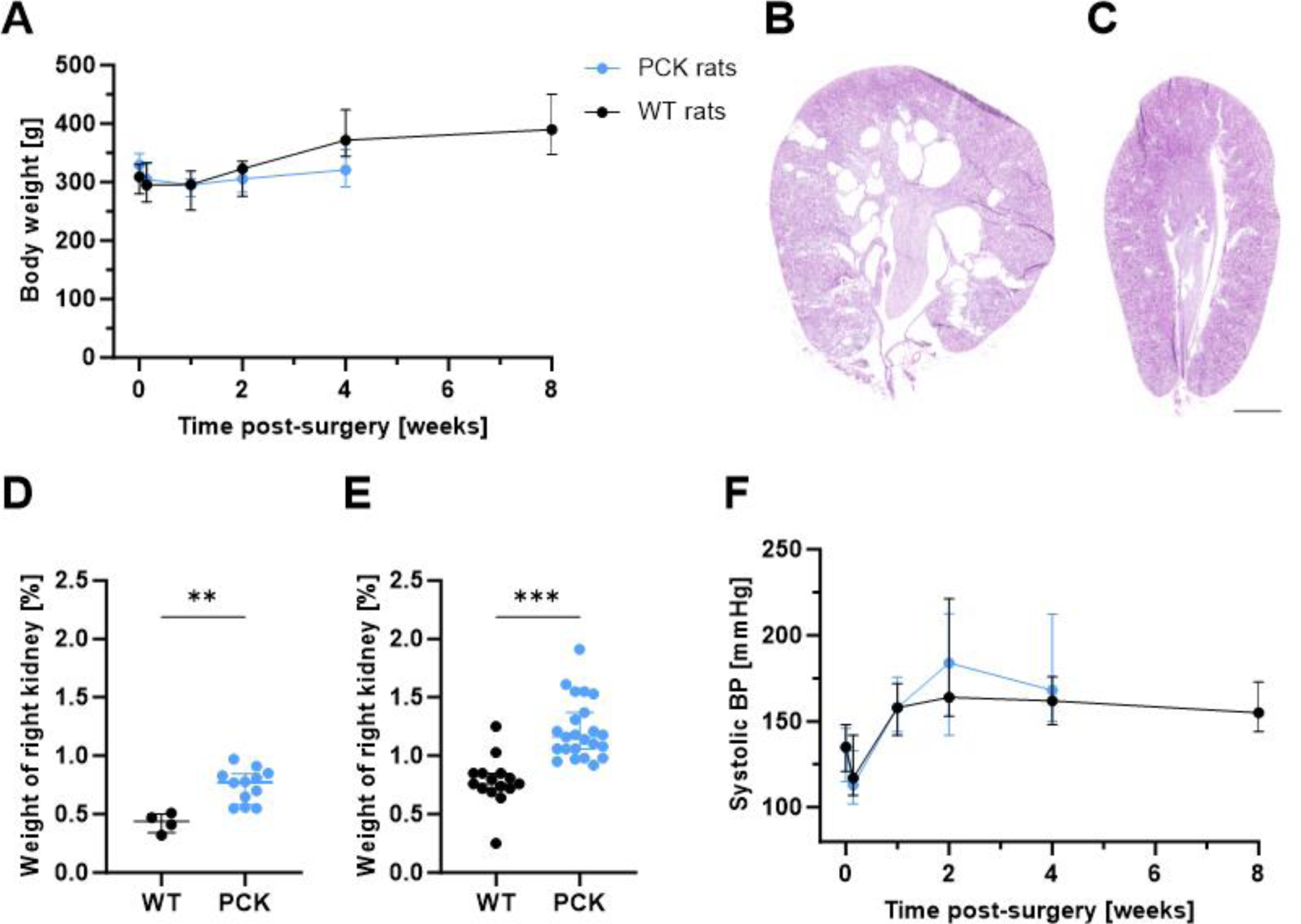
WT and PCK rats have similar bodyweight and blood pressure. **A**, Body weight evolution of WT (n=27) and PCK rats (n=23) following surgery to induce IAs. **B and C**, Hematoxylin/eosin staining of a right kidney section of a PCK (**B**) and a WT (**C**) rat without surgery. Scale bar, 2 mm. **D and E**, Weight of right kidney of control WT (n=4) and PCK (n=12) rats (**D**) and WT (n=15) and PCK (n=23) rats after surgery €. Values are expressed as percentage of body weight. Mann-Whitney U-test. **F**, Systolic blood pressure (BP) evolution of WT (n=27) and PCK (n=23) rats following surgery to induce IAs. **P<0.01, ***P<0.001.

### PCK rats show decreased survival

We originally planned to compare IA development in groups of young WT and PCK rats for up to 8 weeks post-surgery. While the survival rate of WT rats was 100% until 6-weeks post-surgery with thereafter a slight decrease to 90% at 8 weeks, a sudden increase in mortality of PCK rats was observed at 3-weeks post-surgery (**Figure 2A**). The survival rate of PCK rats even dropped below 50% at 4-weeks post-surgery. Thus, in accordance with 3R principles, we decided to stop the experiment with PCK rats at maximum 4-weeks post-surgery. In consequence, WT rats were killed at 2-, 4- and 8-weeks post-surgery, as planned, and the PCK rats at 1-, 2-, 3- and 4-weeks post-surgery. Autopsy of WT and PCK rats that died prematurely during the experiment (3 out of 30 WT and 10 out of 33 PCK rats) revealed the presence of thoracic hemorrhage. Microscopic observation revealed the presence of dissection in the aortic arch (**Figure 2B**) as cause of the thoracic hemorrhage. Disruption of elastic laminae and presence of erythrocytes and fresh fibrin within the arterial wall of the aortic arch, creating a pseudolumen, confirmed the aortic dissection diagnosis (**Figure 2C, 2D and 2E**).

**Figure 2:**
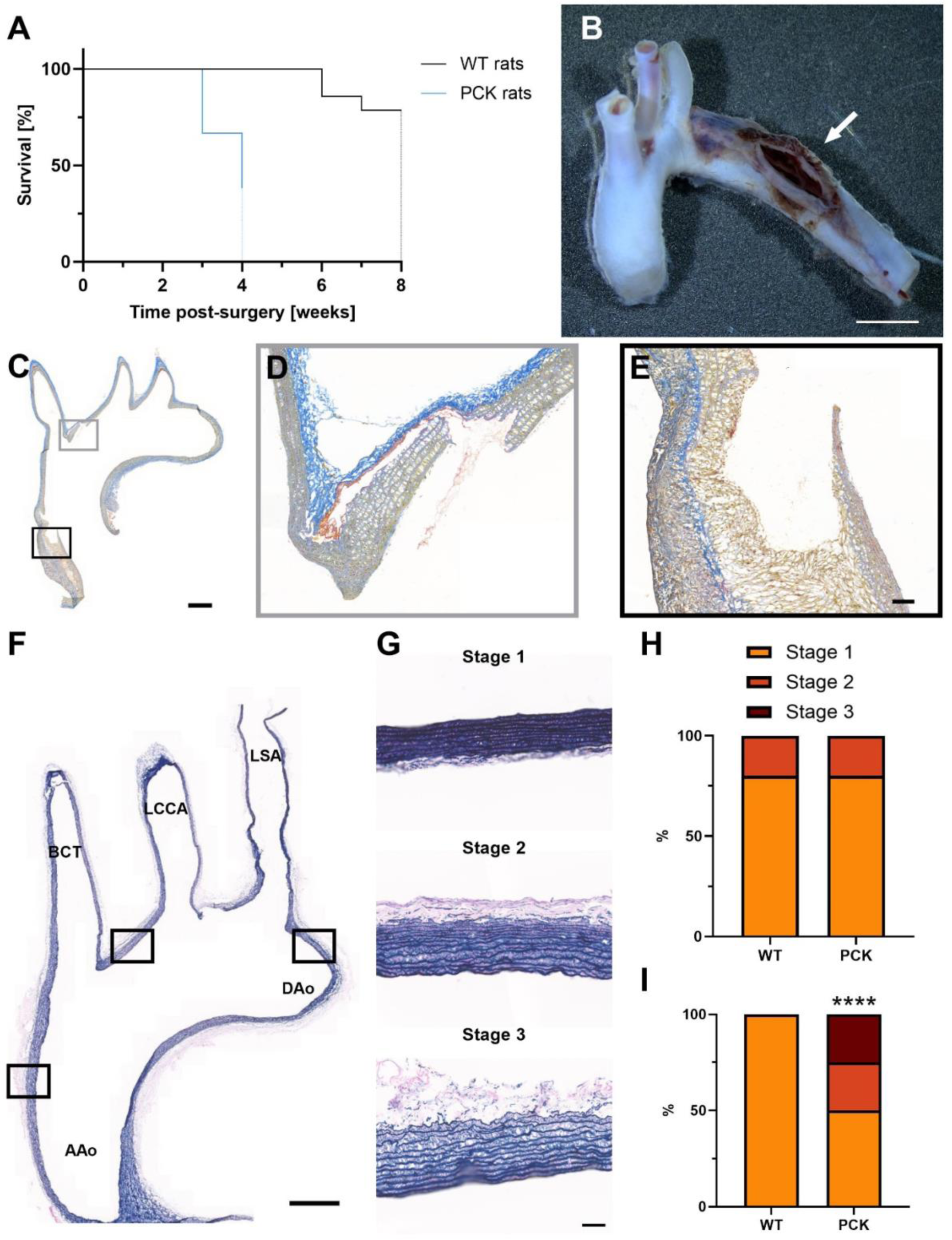
Reduced survival of PCK rats due to aortic dissection. **A**, Survival rate of WT (n=30) and PCK rats (n=33) following surgery to induce IAs. Dotted lines indicate rats killed at the end of the experiment. **B**, Representative image of an aortic arch from a PCK rat that prematurely died 3-weeks post-surgery. Scale bar, 3 mm. Arrow indicates the site of dissection. **C**, Martius Scarlet blue staining of an aortic arch section of a WT rat that prematurely died 6-weeks post-surgery. Nuclei are stained in brown, fibrin in red, erythrocytes and fresh fibrin in yellow, and white blood cells and collagen in blue. Scale bar, 1 mm. Grey (**D**) and black (**E**) panels are magnifications showing the vessel wall deterioration and the infiltration of blood into the vessel wall. Scale bar, 0.1 mm. **F**, Victoria Blue staining of an aortic arch section from a PCK rat killed 4-weeks post-surgery illustrating the 3 locations at which elastic architecture was studied. Scale bar, 1 mm. Aao: ascending aorta, DAo: descending aorta, BCT: brachiocephalic trunk, LCCA: left common carotid artery, LSA: left subclavian artery. **G**, Representative Victoria blue staining images of stages 1, 2 and 3. **H and I**, Classification of the elastic architecture downstream of the LSA in the aortic arch (location indicated in panel D) of 5 control WT and 5 control PCK rats (**H**) and 5 WT and 4 PCK rats killed 4-weeks post-surgery (**I**) Blue, stage 1; red, stage 2; and green, stage 3. Fisher’s exact test. ****P<0.0001

To better understand the aortic wall changes leading to the aortic dissection, the elastic architecture was studied at different locations in the aortic arch (**Figure 2F**) of 5 control WT and PCK rats and 5 WT and 4 PCK rats killed 4-weeks post-surgery. We observed 3 types of elastic architecture (**Figure 2G**): Aortic walls containing straight, thick elastic lamellae and dense elastic fibers were classified as stage 1. Aortic walls containing straight, thick elastic lamellae and not very dense elastic fibers were classified as stage 2. Aortic walls containing thin or undulated elastic lamellae and sparse elastic fibers were classified as stage 3. While the elastic architecture downstream of the left subclavian artery (LSA) was identical and mostly normal (80% stage 1) in the WT and PCK rats without surgery (**Figure 2H**), the elastic architecture was more severely affected at the same location at 4-weeks post-surgery in PCK rats compared to WT rats (**Figure 2I**). This different response of PCK rats to the IA induction protocol as compared to WT rats was even more evident at the level of the ascending aorta and in the aortic wall between the bifurcations of the brachiocephalic trunk (BCT) and the LCCA with elastic architecture classified as stage 2 or 3 in all PCK rats at 4-weeks post-surgery (**Figure S2**). The severe alteration of the elastic architecture in the aortic arch of PCK rats compared to WT rats might explain the increased frequency of aortic dissections observed in PCK rats 4-weeks post-surgery.

### PCK rats display faster IA development

The right OA-ACA bifurcation from WT (**Figure 3A and 3B**) and PCK rats (**Figure 3C and 3D**) were closely examined to find IAs at the various time points post-surgery. Meticulous observation under the dissection microscope revealed the presence of bulging in WT and PCK rats **(Figure 3A and 3C**). Immunofluorescent staining confirmed the bulging and revealed the presence of internal elastic lamina disruption **(Figure 3B and 3D**), which is a characteristic of IAs. WT rats showed a progressive increase in IA occurrence with time after surgery leading to 82% of rats having an IA at 8-weeks post-surgery (**Figure 3E**). Interestingly, PCK rats displayed at 2-weeks post-surgery enhanced IA development (**Figure 3F**) and the maximal IA induction of 86% was already reached at 3-weeks post-surgery in PCK rats, half of the time needed for WT rats (**Figure 3G**). Thus, IA development was faster in PCK rats, which may be due to the absence of functional primary cilia to dampen the endothelial response to disturbed blood flow.

**Figure 3:**
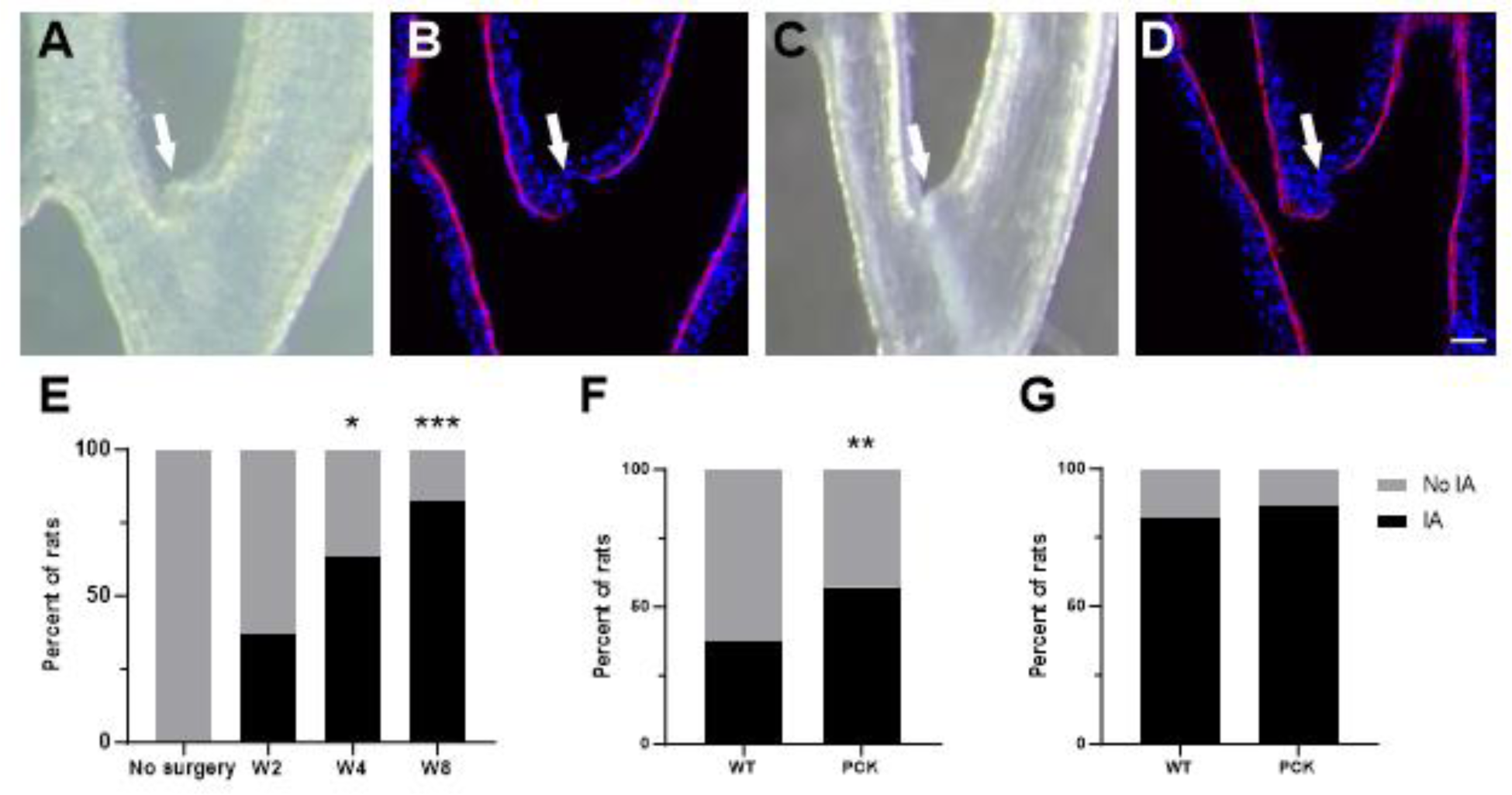
Faster IA development in PCK rats. **A through D**, Representative images of IAs (white arrow) in a WT rat killed 8-weeks post-surgery (**A and B**) and a PCK rat killed 4-weeks post-surgery (**C and D**) observed under the dissection microscope (**A and C**) and sectioned and stained with DAPI (nuclei in blue) and Evans blue (elastic lamina in red) (**B and D**). Scale bar, 50 μm. **E**, Quantification of IA development in WT rats over the 8-weeks experiment (n=8-11/group). Fisher’s exact test with controls without surgery. **F**, Comparison of IA development between 8 WT and 7 PCK rats 2-weeks post-surgery. Fisher’s exact test. **G**, Comparison of IA development between 11 WT rats killed at 8-weeks post-surgery and 7 PCK rats killed at 3-weeks post-surgery. *P<0.05, **P<0.01, ***P<0.001.

### Anatomical variations in the CoW of PCK rats and PKD patients

Disturbed blood flow patterns play a critical role in the initiation of IAs^5^. Anatomical variations in the CoW have been associated with IA formation^9,10^, however the role of primary cilia in this process remains to be investigated. Three frequent variations in the organization of the CoW were observed in the rat, i.e. an additional MCA bifurcation (**Figure 4A**), an additional ACA bifurcation (**Figure 4B**) and the presence of an ACOM (**Figure 4C**). These variants were more frequently observed in PCK rats than in WT rats **(Figure 4D, 4E and 4F**). Therefore, we decided to compare the anatomical variability of the CoW between PKD and non-PKD patients. The @neurIST database contained MRIs of 16 PKD patients. Sixteen non-PKD patients were selected from the same database based on matching for sex, positive family history for IAs and IAs multiplicity. As presented in **Table 1**, this clinical data from PKD and non-PKD patients were not different. Human CoW were classified using MRIs (**Figure S1**) and differences were separated for the anterior (**Figure 4G**) and posterior (**Figure 4H**) part of the CoW. Anterior and posterior CoW of PKD patients were less often classified as type A which corresponds to the predominant CoW anatomical morphology in non-PKD patients. Additional ACOM or ACA were more frequently observed in the anterior part of the CoW from PKD patients than non-PKD patients. Absence or hypoplasia of the ACA was observed in none of the PKD patients while this variant was observed in 12.5% of non-PKD patients. Dominance of one PCOM was observed less frequently and absence or hypoplasia of both PCOM was observed more frequently in PKD patients than in non-PKD patients. Altogether, we observed an increased number of anatomical variations in CoW of both PCK rats and PKD patients, suggesting that the absence of functional endothelial primary cilia might lead to altered embryologic development of the circulatory system.

**Figure 4:**
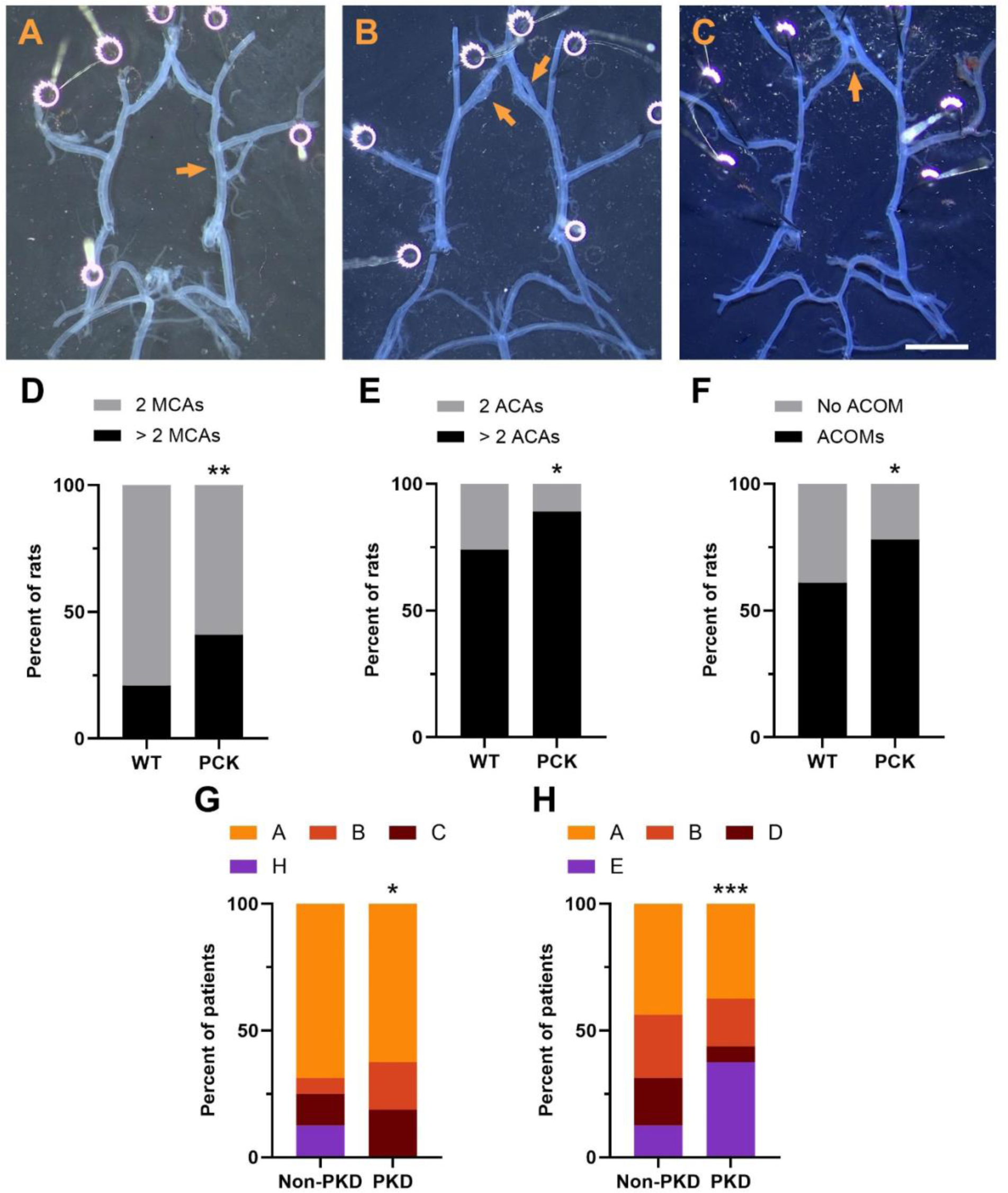
More variation in circle of Willis of PCK rats and PKD patients. **A through C**, Representative images of three variations in the CoW observed in WT and PCK rats (arrows): additional MCA bifurcation (**A**), additional ACA bifurcation (**B**) and presence of an ACOM (**C**). Scale bar, 2 mm. MCA, middle cerebral artery. ACA, anterior cerebral artery. ACOM, anterior communicating artery. **D though F**, Corresponding quantifications of these three CoW variations in WT (n=38) and PCK (n=46) rats. Fisher’s exact test. **G and H**, Prevalence of anterior (**G**) and posterior (**H**) variants of the CoW of PKD (n=16) and matched non-PKD (n=16) patients. Fisher’s exact test. *P<0.05, **P<0.01, ***P<0.001.

**Table 1.** Clinical data of the PKD and non-PKD patients.

|  | PKD patients | Non-PKD patients | p-value |
| --- | --- | --- | --- |
| Patients | n = 16 | n = 16 |  |
| Age, year: mean (SD) | 55 (16) | 64 (13) | 0.09 |
| Sex, female: n (%) | 10 (63) | 13 (81) | 0.43 |
| Multiple **IAs**: n (%) | 8 (50) | 7 (44) | >0.99 |
| Previous aSAH: n (%) | 0 (0) | 0 (0) | >0.99 |
| Positive family history for IA: n (%) | 5 (31) | 3 (19) | 0.69 |
| Smoker (former and current): n (%) | 9 (56) | 8 (50) | >0.99 |
| Hypertension: n (%) | 12 (75) | 8 (50) | 0.27 |

Comparison of means have been performed using two-tailed Student's *t*-test. Comparison of distribution have been performed using Fisher's exact test. SD, standard deviation. aSAH: aneurysmal subarachnoid hemorrhage. IA: intracranial aneurysm.

### Absence of increase in tight junction proteins in IAs of PCK rats

In an unbiased transcriptomics *in vitro* study, we previously identified ZO-1 as a central regulator of primary cilia-dependent TJ integrity and endothelial permeability^13^. To investigate the relation between primary cilia and endothelial junction integrity *in vivo*, we examined different EC junctional proteins in IAs at the OA-ACA bifurcation and along the ACA, a region free of IA development, in control groups of WT and PCK rats and at 2- and 4-weeks post-surgery (**Figure 5A, 5B, 5C and 5D**). Importantly, staining for the endothelial marker CD31 was similar in all groups of WT and PCK rats both at the level of the ACA and OA-ACA bifurcation (**Figures 5B, 5E and 5F**), illustrating that the presence of endothelium was similar at both locations and not affected by the surgical procedure in rats with and without functional primary cilia. Next, we focused more specifically on the expression of the TJ proteins ZO-1 (**Figure 5C**) and CLDN5 (**Figure 5D**) in the endothelium. At the level of the ACA, a similar increase of ZO-1 staining was observed 2- and 4-weeks post-surgery in ECs from WT and PCK rats in comparison to WT and PCK rats without surgery (**Figure 5G**). At the level of the IA, the ZO-1 staining was greatly increased in WT rats 2-weeks post-surgery whereas this increase was absent in PCK rats (**Figure 5H**). Likewise, CLDN5 was increased in the OA-ACA bifurcation of WT rats, and not of PCK rats, at 2- and 4-weeks post-surgery (**Figure 5J**).

**Figure 5:**
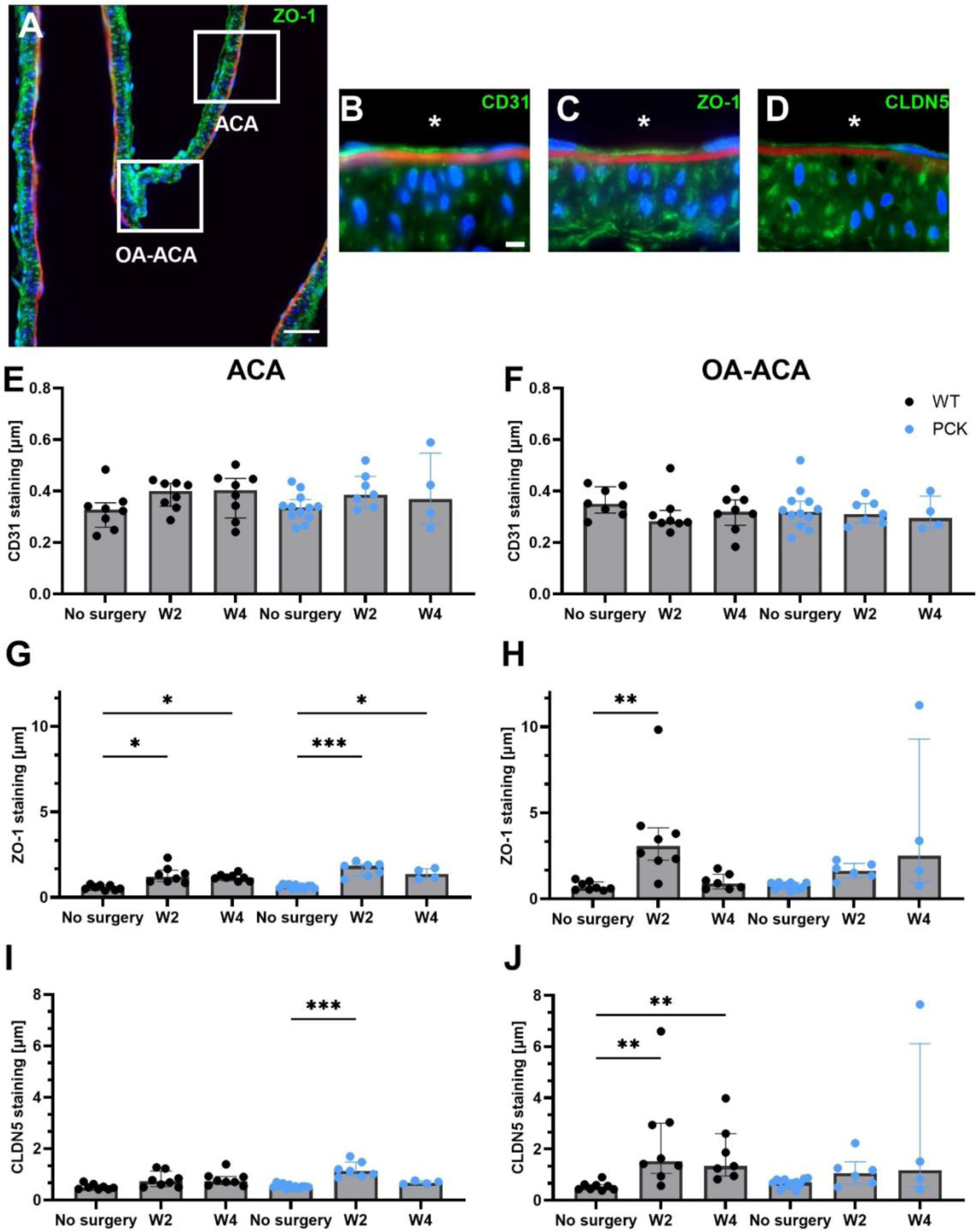
Absence of increase in endothelial TJ proteins in IAs of PCK rats. **A**, Representative ZO-1 immunostaining of the right OA-ACA bifurcation of a WT rat killed 8-weeks post-surgery. The white squares indicate two zones of interest: the OA-ACA bifurcation where IAs develop and a straight part of the ACA. Scale bar, 50 μm. **B thought D**, Representative CD31 (**B**), ZO-1 (**C**) and CLDN5 (**D**) immunostainings (in green). Nuclei are stained with DAPI (in blue), and internal elastic laminae are stained with Evans blue (in red). * denotes lumen of the vessel. Scale bar, 5 μm. **E through J**, Quantifications of CD31 (**E and F**), ZO-1 (**G and H**) and CLDN5 (**I and J**) immunostainings on WT (n=7-8/group) and PCK (n=4-11/group) rats killed without surgery, at 2-weeks and 4-weeks post-surgery in the ACA (**E, G and I**) and at the OA-ACA bifurcation (**F, H and J**). Kruskal-Wallis test with Dunn’s multiple comparison test with WT and PCK without surgery. *P<0.05, **P<0.01, ***P<0.001.

Finally, the same endothelial TJ proteins were studied in 19 vulnerable (ruptured) and 38 more stable (unruptured) human IA domes. Clinical data from patients with ruptured and unruptured IAs were not different (**Table 2**). As expected, maximal IA diameter and bottleneck factor were larger in ruptured IAs compared to unruptured IAs. Alike the rat IAs, ZO-1 immunostaining was lower in ruptured IAs compared to unruptured IAs (**Figure 6A-6C**). A similar decrease was observed for CLDN5 in ruptured IAs (**Figure 6D-6F**), while no difference in CD31 staining was observed between ruptured and unruptured IAs (**Figure 6G-6I**). The decrease of TJ proteins observed in IAs from PCK rats and in vulnerable human IAs might explain why IA in PKD patients at more at risk to rupture.

**Figure 6:**
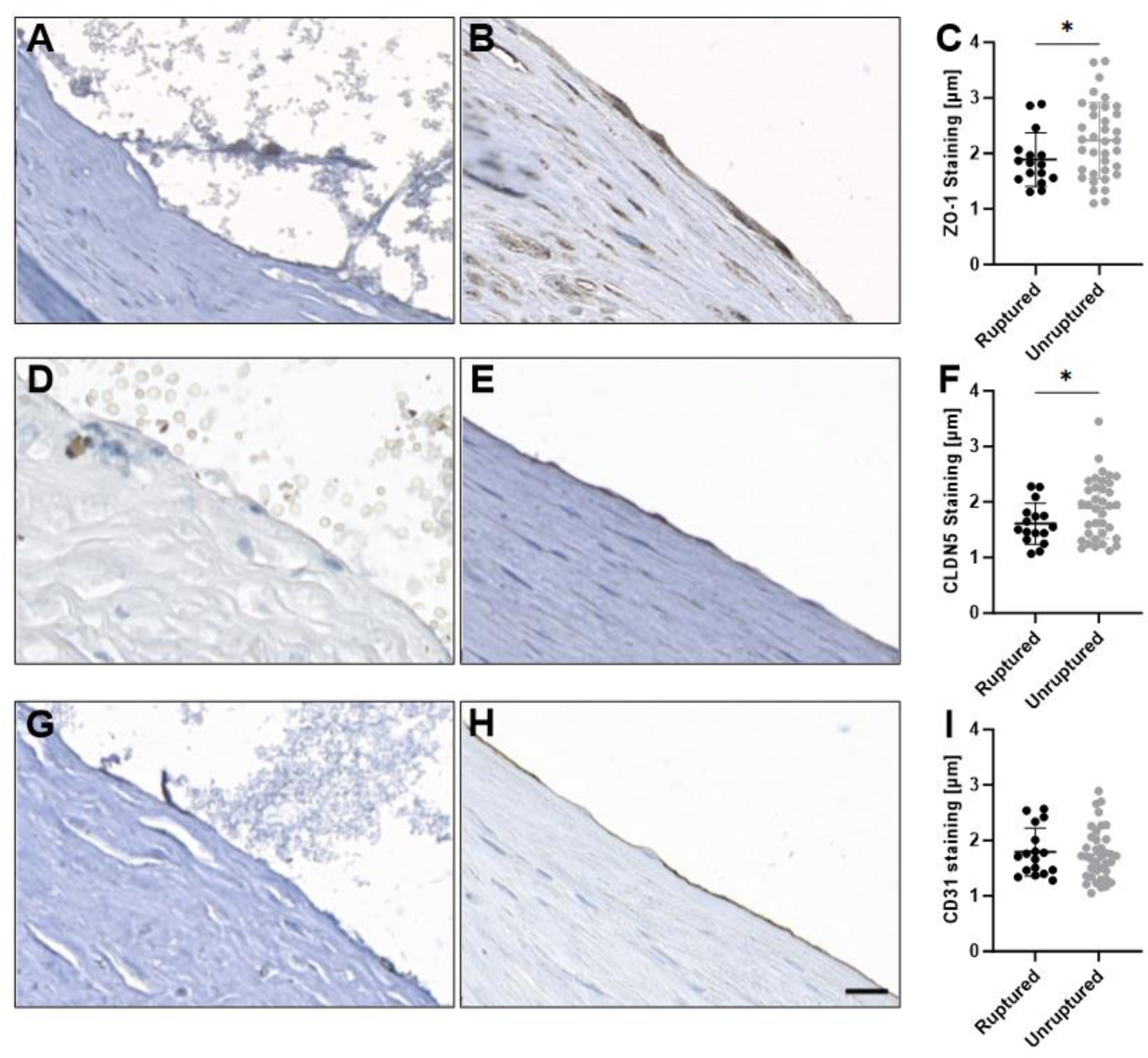
Decreased endothelial TJ proteins in ruptured IAs of patients. **A through C**, ZO-1 immunostaining (in brown) on ruptured (**A**) and unruptured (**B**) human IAs and corresponding quantification (**C**) on ruptured (n=16) and unruptured (n=36) IAs. **D through F**, CLDN5 immunostaining (in brown) on ruptured (**D**) and unruptured (**E**) human IAs and corresponding quantification (**F**) on ruptured (n=16) and unruptured (n=36) IAs. **G through I**, CD31 immunostaining (in brown) on ruptured (**G**) and unruptured (**H**) human IAs and corresponding quantification (**I**) on ruptured (n=17) and unruptured (n=38) IAs. One-tailed Student’s *t*-test. Scale bar, 20 μm. *P<0.05.

**Table 2.** Patient and IA characteristics of patients with ruptured and unruptured IAs.

|  | Ruptured IAs | Unruptured IAs | p-value |
| --- | --- | --- | --- |
| **Patients** | **n = 19** | **n = 38** |  |
| Age, year: mean (SD) | 51 (11) | 56 (11) | 0.14 |
| Sex, female: n (%) | 14 (74) | 27 (71) | >0.99 |
| Multiple **IAs**: n (%) | 8 (44) <sup>▲</sup> | 20 (53) | 0.78 |
| Previous aSAH: n (%) | 2 (11) <sup>▲</sup> | 5 (13) | >0.99 |
| Positive family history for IA: n (%) | 2 (11) <sup>▲</sup> | 5 (14) <sup>▲</sup> | >0.99 |
| Smoker (former and current): n (%) | 12 (67) <sup>▲</sup> | 28 (74) | 0.75 |
| Hypertension: n (%) | 6 (33) <sup>▲</sup> | 14 (37) | >0.99 |
| **Intracranial aneurysm** | **n = 19** | **n = 38** |  |
| Rough appearance: n (%) | 5 (33) <sup>▼</sup> | 17 (46) <sup>▲</sup> | 0.54 |
| Presence of blebs or lobules: n (%) | 13 (72) <sup>▲</sup> | 20 (53) | 0.25 |
| Neck size, mm: median (IQR) | 3.6 (2.6-4.9) <sup>►</sup> | 3.8 (3.0-5.0) | 0.79 |
| Maximal diameter, mm: median (IQR) | 7.9 (5.8-10.6) <sup>▲</sup> | 5.7 (4.2-8.2) | 0.03 |
| Bottleneck factor: median (IQR) | 2.0 (1.4-2.7) <sup>►</sup> | 1.6 (1.2-1.8) | 0.03 |

Comparison of means and medians have been performed using two-tailed Student's *t*-test and Mann- Whitney U-test, respectively. Comparison of distribution have been performed using Fisher's exact test. SD, standard deviation. IQR, interquartile range. aSAH: aneurysmal subarachnoid hemorrhage. IA: intracranial aneurysm. Missing information for 1 (<sup>▲</sup>), 3 (<sup>►</sup>) or 4 (<sup>▼</sup>) patients.

## DISCUSSION

The mechanisms leading to increased risk for PKD patients to develop more IAs of increased vulnerability are still not completely understood. In this study, we used young PCK rats that have a similar weight curve than WT rats (until maximal 14 weeks of age) but develop cysts in the kidneys already at 21 days of age^19^. Despite the presence of cysts in the kidneys, PCK rats did not develop a higher blood pressure than WT rats following the protocol to induce IAs **(Figure 1**). This protocol however drastically increased the mortality of PCK rats after 3-weeks post-surgery. We uncovered that aortic dissection causing fatal thoracic hemorrhage was the cause of death (**Figure 2**). Interestingly, IAs developed faster in PCK rats compared to WT rats (**Figure 3**), suggesting that PCK rats are more sensitive to IA induction. In addition, we observed an increased frequency of anatomical variations in the CoW of PCK rats and PKD patients compared to WT rats and non-PKD patients (**Figure 4**). Finally, the increased expression of TJ proteins observed in WT rats after IA induction was not observed in PCK rats (**Figure 5**). A similar decrease in TJ protein expression was observed in vulnerable (ruptured) IAs compared to more stable (unruptured) IAs (**Figure 6**). Thus, variations in the anatomy of the CoW and impaired regulation of TJ proteins might put PCK rats and PKD patients more at risk to develop vulnerable IAs.

The protocol to €nduce €As unexpectedly increased the mortality of PCK rats. It has been previously described that PKD symptoms affect PCK rats and decrease their survival after 17 weeks of age^18^. In our experiment, we observed a decreased survival at 3-weeks post-surgery, which corresponds to 9- to 13-weeks old rats. It is thus unlikely that the premature death of PCK animals in our study resulted directly from PKD and we assume that the aortic dissection resulting in the premature death of PCK rats is a consequence of the IA induction protocol. Of further note, the increased frequency of aortic dissection is also unlikely to be a direct consequence of PKD-induced hypertension since we observed similar levels of hypertension in WT and PCK rats. Angiolathyrism induced by the lathyritic compound BAPN, which inhibits the cross-linking between elastin and collagen^23^, facilitates IA development in combination with increased hemodynamic stress and hypertension in this IA rat model^24^. BAPN affects not only intracranial arteries, but all blood vessels and thoracic hemorrhage has been described by Hashimoto and colleagues in WT rats exposed to the IA induction protocol^25^. In this study, however, BAPN was described to mostly affect young and growing animals and to have less effect in adult WT rats. Our study revealed severe alterations in the elastic architecture of the aortic wall in adult PCK rats at 4-weeks post-surgery, to which BAPN certainly contributed. Pulsatile cardiac outflow at high pressure might be an additional damaging factor given the observed more severe morphological alterations in the ascending aorta. It is noteworthy that PKD patients are not only at higher risk of IA development but also display an increased risk of developing aortic aneurysms and dissections^26,27^. Altogether, we postulate that the lack of functional primary cilia in the endothelium of PCK rats in combination with the induced hypertension and the BAPN treatment drastically increased the risk of developing aortic dissection €n the PCK rats through an alteration of the elastic architecture.

A majority of the population lacks a full CoW^28^ and variations in the CoW have been correlated with IA disease^9,10^. Moreover, the CoW of women, who are at greater risk to develop IAs, was shown to be different from men and the distribution of IA locations was also shown to be different^29,30^. In our study, we observed more anatomical variation in the CoW of PKD patients and PCK rats than in the CoW of non-PKD patients and WT rats. Furthermore, a recent meta-analysis onto IA distribution in patients with autosomal dominant PKD revealed different IA locations in this subgroup than in the general population^31^. Indeed, PKD patients developed more often IAs in large diameter arteries like MCA, ICA and ACOM^31^. As a different organization modifies the blood flow pattern within the CoW, the different IA locations in PKD patients might be due to their different CoW organization, as suggested by the results of this study. The embryonic development of the CoW is greatly influenced by genetic factors^28,32^. Moreover, the development of the vascular system during embryogenesis is highly regulated by blood flow^33^. Therefore, lack of functional primary cilia in Ecs of PKD patients might impact the arterial growth and branching during embryogenesis leading to a different CoW organization. Subsequently, the variation €n hemodynamics resulting from this different CoW organization explains the increased risk to develop IAs. In addition, variability of the CoW configuration has been associated with an increased risk of IA rupture^34,35^. Thus, screening for anatomical variations in the CoW might help to identify non-PKD patients that are at greater risk of IA rupture and may further open the way to personalized decisions for IA treatment.

In accordance with an earlier study in different mouse models of PKD^36^, we found that IAs developed faster in PCK rats compared to WT rats. As this rat and the mouse PKD model carry different mutations, this strongly suggests a causal relation between the absence of functional endothelial primary cilia and the sensitivity to develop IAs. As previously demonstrated *in vitro*, primary cilia dampen the endothelial response to an aneurysmal pattern of disturbed flow^13^. Thus, an exaggerated endothelial response to disturbed blood flow may cause more IAs in PKD patients. Interestingly, PKD patients have not only an increased sensitivity to IA development but they are also classified as a high risk group for aneurysmal SAH^37^, suggesting that their IA are more susceptible to rupture.

We previously observed a reduction of the TJ protein ZO-1 in the endothelium of IA domes from PKD patients in comparison to non-PKD patients^13^. As ZO-1 is crucial for an intact endothelial barrier, we postulated that a decrease of ZO-1 might explain why IAs of PKD patients are more at risk of rupture. To investigate a possible causal relation between the absence of functional endothelial primary cilia and IA vulnerability, we investigated PCK rats and studied the expression of the TJ proteins ZO-1 and CLDN5 in the endothelium of IAs at the OA-ACA bifurcation, a preference location of primary cilia^13^, and in the endothelium of a non-IA region of the ACA. Ecs in WT and PCK rats showed a similar increase in ZO-1 and CLDN5 staining at the level of the ACA 2- and 4-weeks post-surgery, which is likely an adaptive response to the increased hemodynamic stress and hypertension. The molecular mechanism responsible for this adaptive response was not studied. However, it is known that endothelial TJ protein expression can be regulated through the Akt1-FoxO signaling pathway^38^ or the Angiopoietin-Tie signaling pathway^39^. Interestingly, at the level of the OA-ACA bifurcation, where IAs develop, the adaptive increase in ZO-1 and CLDN5 staining was only observed in WT rats. The absence of this protective effect at the OA-ACA bifurcation might be a further explanation as to why IAs develop faster in PCK rats. In addition, the lack of an adaptive TJ-strengthening response in PCK rats may also facilitate macrophage infiltration over the endothelial barrier into the IA as postulated by Tada *et al*.^40^, thereby enhancing inflammation and vulnerability of the IA.

A limitation of our study is that only 4 PCK rats survived up to 4-weeks post-surgery making it less reliable to draw conclusions on IA development in PCK rats after 3-weeks post-surgery. In addition, the IA rat model used in this study gives rise to stable IAs with a very low rupture rate in WT rats^41^. Although IAs in PCK rats seemed to have a more vulnerable phenotype based on TJ protein expression, our study did not investigate IA rupture and can thus only postulate towards the increased risk of IA rupture in PKD patients. Other IA animal models exist to study IA rupture^42,43^, but PCK rats might be too sensitive to such protocols given the drastic decrease in survival following the present IA induction protocol.

In conclusion, PCK rats are more vulnerable to IA induction and developed IAs faster than WT rats. IA development is a complex process influenced by several factors. We observed variations in the CoW anatomy and an impaired regulation of TJ proteins in PCK rats which might explain the increased incidence of IAs in PKD patients. Moreover, these differences may also explain why PKD patients are more at risk of IA rupture. Finally, these differences might help to identify non-PKD patients who are at greater risk of IA rupture.

## Nonstandard Abbreviations and Acronyms

AAo: ascending aorta
ACA: anterior cerebral artery
ACOM: anterior communicating artery
BA: basilar artery
BAPN: β-aminopropionitrile
BCT: brachiocephalic trunk
CCA: common carotid artery
CLDN5: claudin-5
CoW: circle of Willis
DAo: descending aorta
ECs: endothelial cells
IA: intracranial aneurysm
ICA: internal carotid artery
LCCA: left common carotid artery
LSA: left subclavian artery
MCA: middle cerebral artery
MRI: magnetic resonance imaging
OA: olfactory artery
PCA: posterior cerebral artery
PCOM: posterior communicating artery
PKD: polycystic kidney disease
SAH: subarachnoid hemorrhage
TJ: tight junction
TOF: time of flight
WT: wild-type
ZO-1: zonula occludens-1

## Acknowledgments

The authors thank Bernard Foglia, Graziano Pelli and Marc Saulnier-Navetier for technical assistance.

## Sources of Funding

This work was supported by the Fondation Privée des HUG (to B.R.K., E.A., P.B.), the Gottfried und Julia Bangerter-Rhyner-Stiftung (to B.R.K.), the Wolfermann-Nägeli-Stiftung (to B.R.K.) and the Swiss Heart Foundation (to B.R.K., E.A., P.B.).

## Disclosures

None.

## Supplementary figures

**Figure S1:**
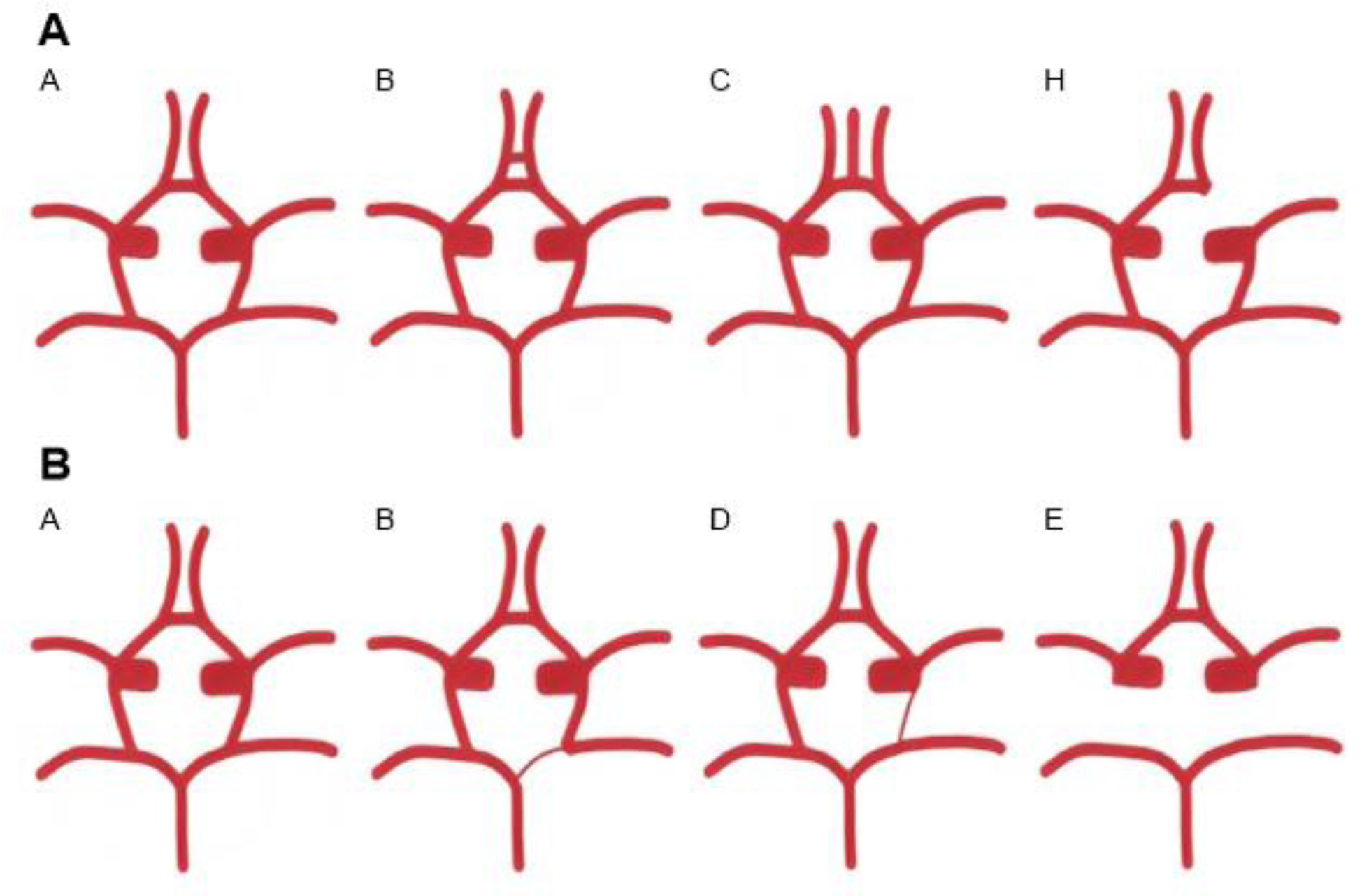
Anatomic variations in the human circle of Willis. **A**, Diagrams of the anatomic variations in the anterior part of the CoW. Variant A: Normal CoW anatomy. Variant B: Additional ACOM(s). Variant C: The median artery of the corpus callosum originates from the ACOM. Variant H: Hypoplasia or absence of the A1 segment of one of the ACAs. **B**, Diagrams of the anatomic variations of the posterior part of the CoW. Variant A: Normal CoW anatomy. Variant B: One PCA originates predominantly from the ICA. This variation is also known as unilateral fetal type PCA. Variant D: Hypoplasia of one PCOM. Variant E: Hypoplasia or absence of both PCOMs. Adapted from Kızılgöz *et al*^6^.

**Figure S2:**
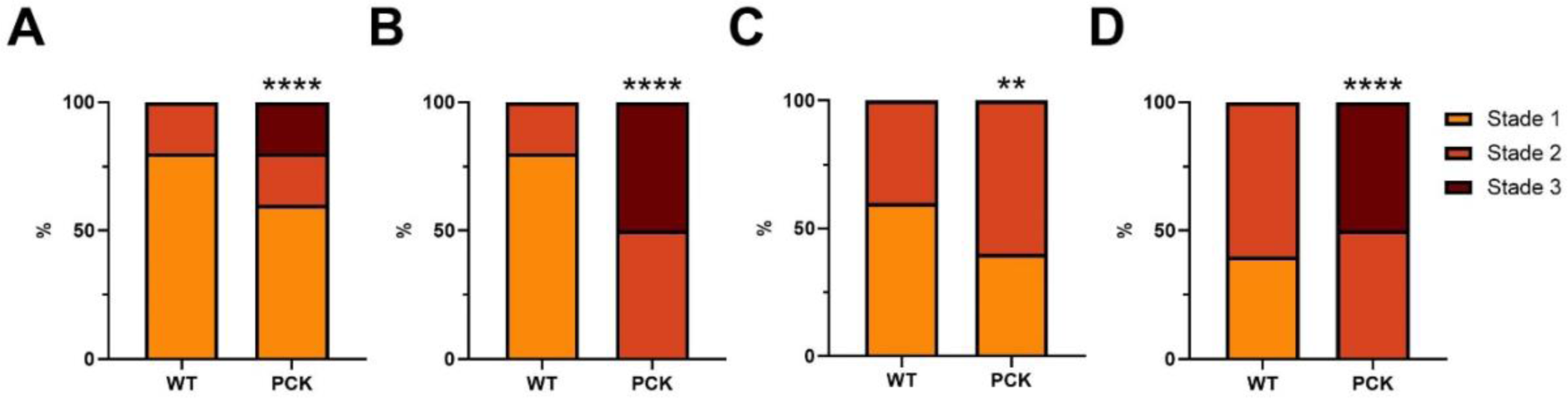
Elastic architecture in the aortic wall of WT and PCK rats. Classification of the elastic architecture in the aortic wall between the bifurcations of the BCT and the LCCA (**A and B**) and in the ascending aorta (**C and D**) (locations indicated in **Figure 2D**) of 5 control WT and 5 control PCK rats (**A and C**) and 5 WT and 4 PCK rats killed 4-weeks post-surgery (**B and D**). Blue, stage 1; red, stage 2; and green, stage 3. Fisher’s exact test.

## Major Resources Table

### Animals

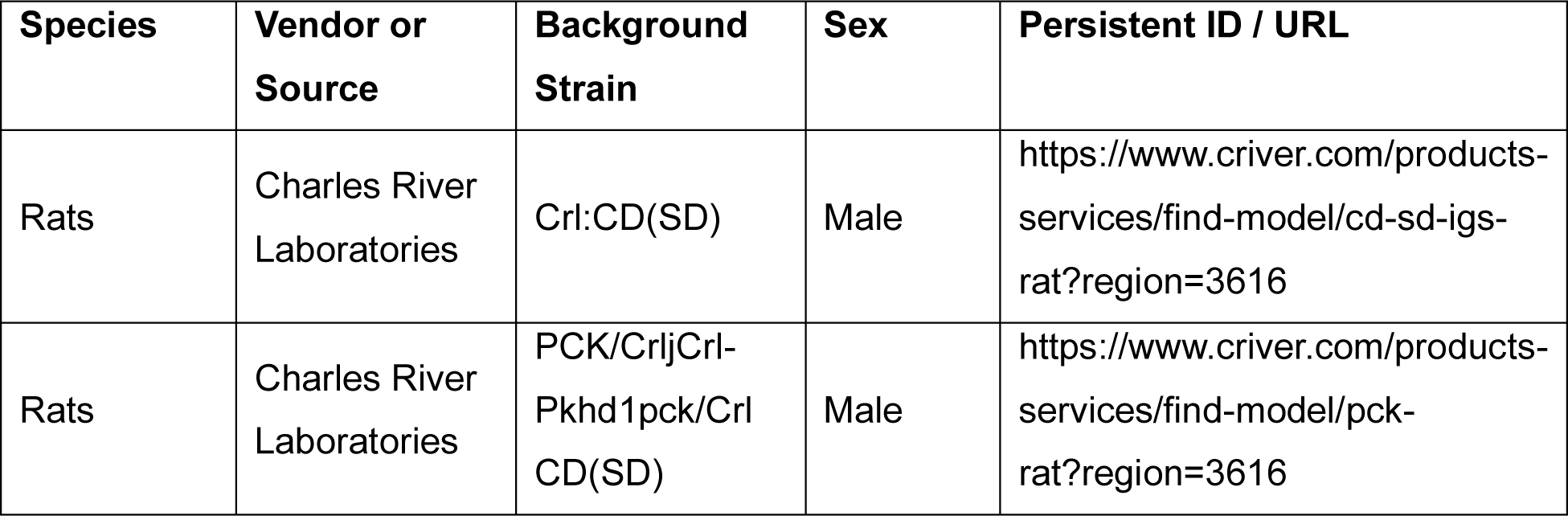

### Antibodies

| Target antigen | Vendor or Source | Catalog # | Working concentration | Persistent ID / URL |
| --- | --- | --- | --- | --- |
| CD31 | Dianova | DLN-33600 | 1/25 (human)<br>1/100 (rat) | https://www.dianova.com/en/shop/dln-033600-anti-cd31-hu-from-rabbit-clone-pab-unconj/ |
| ZO-1 | Invitrogen | 61-7300 | 1/100 | https://www.thermofisher.com/antibody/product/ZO-1-Antibody-Polyclonal/61-7300 |
| CLDN5 | Abnova | PAB27037 | 1/1000 (human)<br>1/200 (rat) | https://www.abnova.com/en-global/product/detail/pab27037 |
| FITC-conjugated goat anti-rabbit | Jackson Lab | 711-095-152 | 1/100 | https://www.jacksonimmuno.com/catalog/products/711-095-152 |

### Chemicals and assay kits

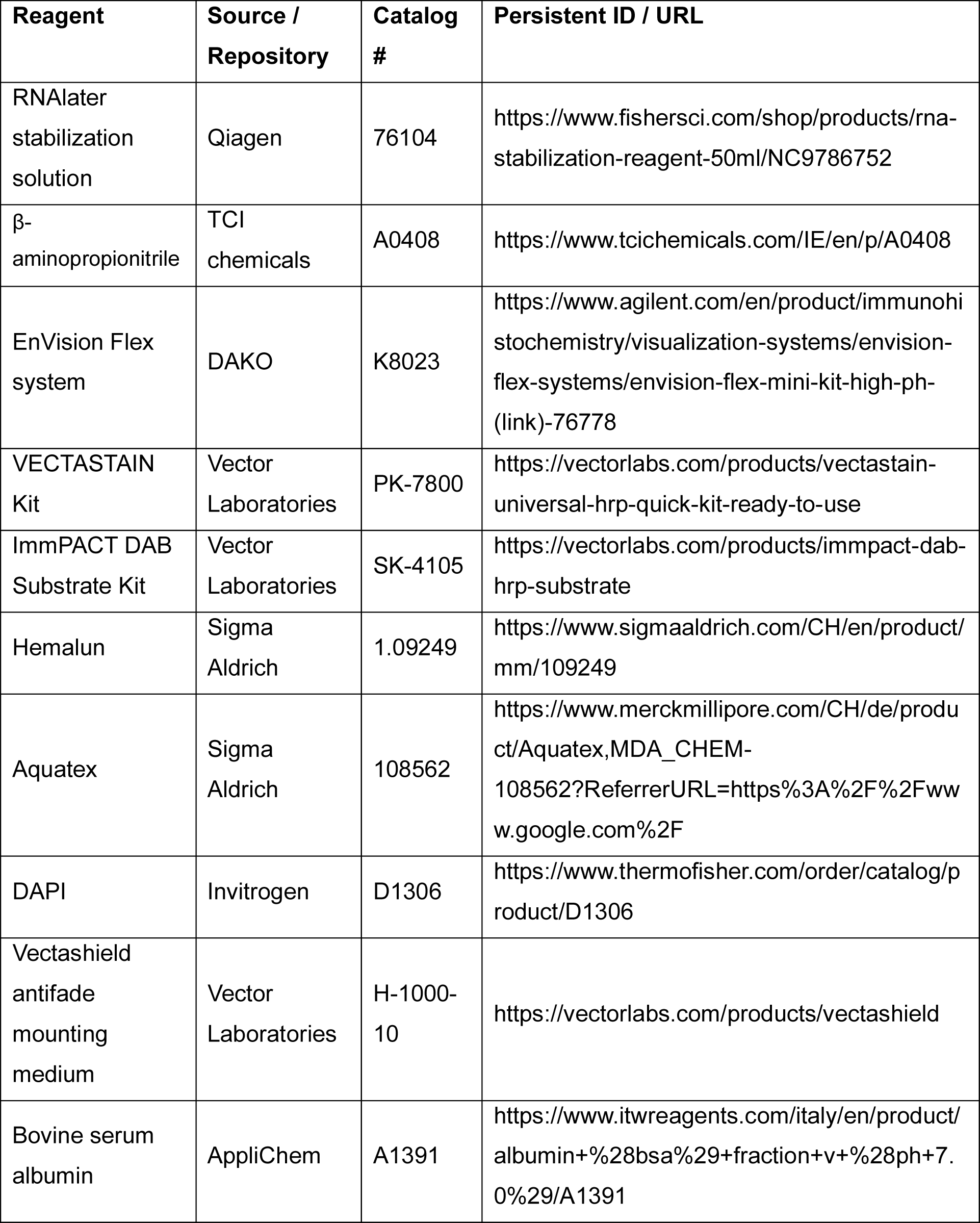

## Notes

### Competing Interest Statement

The authors have declared no competing interest.

